# Equations describing semi-confluent cell growth (I) Analytical approximations

**DOI:** 10.1101/2023.08.27.554985

**Authors:** Damien Hall

## Abstract

A set of differential equations with analytical solutions are presented which can quantitatively account for the effects of variable degrees of contact inhibition on cell growth in two- and three-dimensional cultures. The developed equations can be used for comparative purposes when assessing the extent of contribution of higher order effects, such as culture geometry and nutrient depletion, on mean cell growth rate. These equations also offer experimentalists the opportunity to characterize their cell culture growth experiments using a single reductive parameter.

---

The cell is the basic ‘quanta’ of biology, and as such, achieving a quantitative understanding of how it grows, reproduces and interacts with surrounding cells under both normal and disease states is of fundamental importance **[Anderson et al. 2007; Karolak et al. 2018; Slack and Dale, 2021;Montagud et al. 2021]**. Typically, when cells divide in a stirred-liquid medium they are separated from each other by shear forces produced by mixing **[Ritacco et al. 2018]**. However, when growing and reproducing on or in a non-stirred viscous, or solid, media, any newly formed cells will exist near to their precursors **[Kapałczyńska et al. 2018]**. Many studies have demonstrated how particular spatial relationships between cells in a growing colony can affect an individual cell’s growth pattern with the most well-known being contact inhibition (or local confluence), in which further growth and division is arrested in cells surrounded by other cells **[Martz and Steinberg, 1972; Lee et al. 1995; Ribatti, 2017; Mendonsa et al. 2018]**. Changes in the growth rate of cells within a colony can be assigned to complex factors, such as nutrient depletion **[Burton, 1966; McElwain and Ponzo, 1977; Drasdo and Höhme, 2005; Banwarth-Kuhn et al. 2020; Golden et al. 2022]**, or the release of chemicals by cells at specific positions within the colony **[Bassler and Losick, 2006; Schmitz et al. 2021]**. It is difficult to tease out the exact weight of these contributions when more basic models detailing the arrest of cell growth by local contact inhibition in a colony are not available. Here, we present analytical forms relevant to describing variable degrees of contact inhibition, in colonies growing in 2D (a circle), and in 3D (a sphere).

## Model development

The division of a cell, C, to yield two cells (a mother and a daughter), is considered to be regulated by a first order rate constant, k (Eqn. 1) **[Hartwell and Unger, 1977; Drasdo and Höhme, 2005]**.

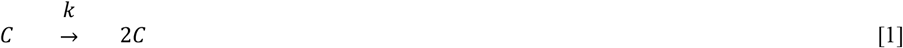

For the case of zero local confluence (no contact inhibition)^1^ the rate of formation of the number of cells, N, at a specific time, t, can be described by a first order differential equation (Eqn. 2) which itself can be integrated to yield the equation for exponential cellular growth (Eqn. 3).

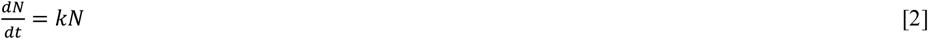

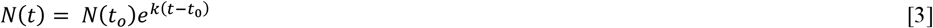

In this work we consider contact inhibition to be simply defined by the immediate local density of cells surrounding the cell of interest^2^ (**Fig. 1a**). Within a colony we may define an outer perimeter region of one cell thickness, hereon described as the colony ‘edge’, which contains cells not subject to contact inhibition and hence able to grow (**Fig. 1b**). Conversely, within the interior region there exist cells, surrounded by other cells, which are subject to some variable degree of contact inhibition defined by the parameter, β, such that when β = 0 there is zero contact inhibition, and when β = 1 there is a complete contact inhibition and the cells are unable to divide **[Lee et al. 1995; Hall, 2023]**. To simplify the subsequent mathematics, we set a complementary transition parameter, γ, that is reflective to β about the point 0.5 (Eqn. 4a). Described in such a way we can define a modified rate constant for cellular growth that accommodates contact inhibition (Eqn. 4b) and can be used in formulating the differential equation for growth (Eqn. 2).

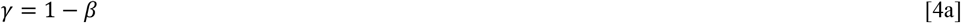

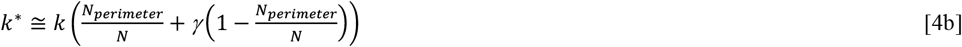

**Figure 1:**
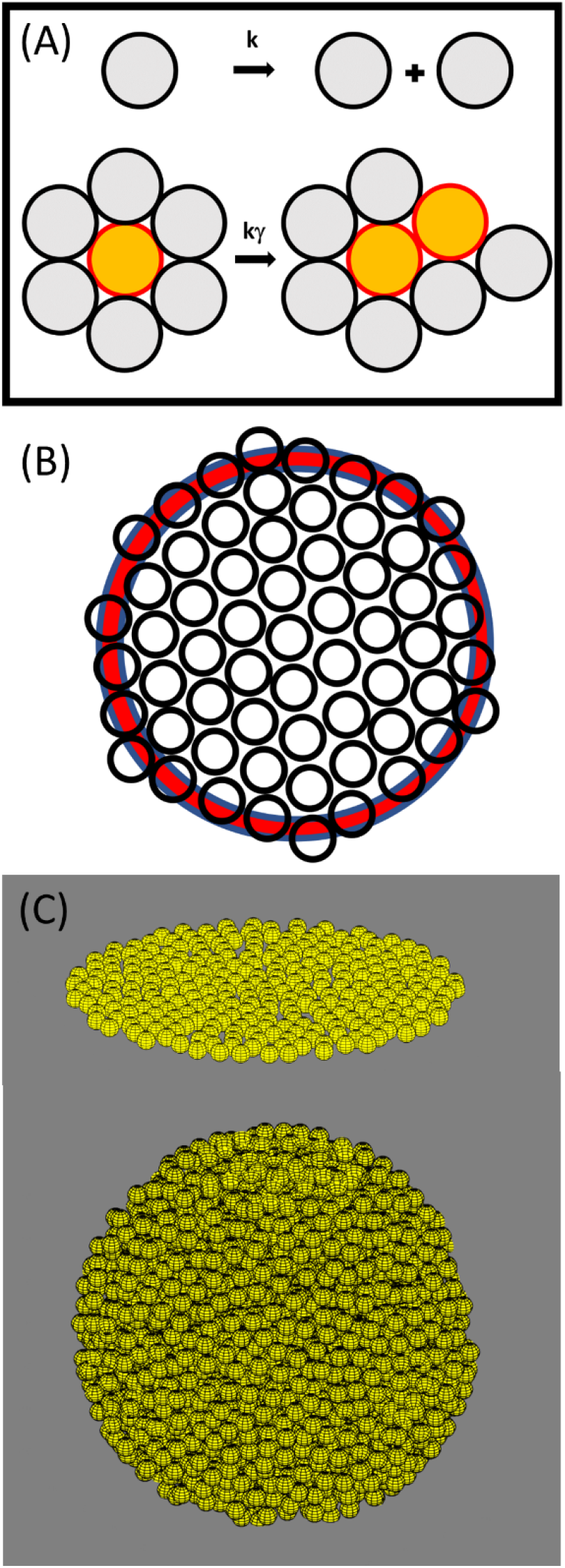
Considerations of cell growth. (A) Cell growth kinetics and local contact inhibition: Cell division and growth is regulated by a first-order rate constant k. Local contact inhibition is considered to occur when cells are surrounded by other cells leading to a reduction in growth rate defined by the parameter γ (Eqn. 4a). (B) Position dependent growth: Colonies of cells are considered in terms of perimeter regions (shown in red) and interior regions (shown in white) (Eqn. 4b). (C) Colony Geometry: Within the present work cells are considered able to grow in one of two types of geometry, a circular 2D monolayer (Eqns 6-8), and a 3D spherical body (Eqns. 9-11).

In keeping with experimental observations of cells grown in two and three-dimensional cultures we consider colony growth in one of two basic geometries, a circular pattern of one cell thickness (cells growing as a monolayer on a flat surface **[Martz and Steinberg, 1972; Wakita et al. 1994; Schnyder et al. 2021]**), and a sphere **[Alessandri et al. 2013; Montel et al. 2012; Karolak et al. 2018; Schmitz et al. 2021]** (**Fig.1c**). To describe the number of cells in the perimeter and interior regions for each of the three culture geometries considered, we first require a description of the state of area/volume occupancy of cells within the colony. To achieve this, we introduce the concept of a packing constant, F, which describes the imperfect filling of the area, or volume, of cells within the culture, such that F ≥ 1 (Eqn. 5).

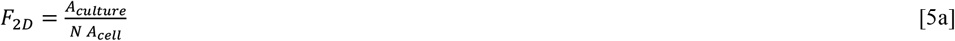

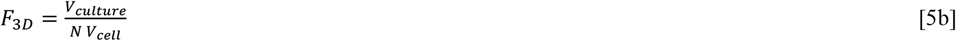

To a first approximation we consider the cells to be spherical and to be characterized by a radius, r_1_. In two-dimensions, values of F_2D_ for the random close packing (RCP) and hexagonal close packing (HCP) of circles within a plane are ∼1.25 and 1.1 **[Kausch et al. 1971]**. In three-dimensions, the corresponding values for F_3D_ in the RCP and HCP cases are respectively ∼1.9 and 1.3 **[Scott and Kilgour, 1969; Lubachevsky et al. 1991]**. We also acknowledge that the imperfect packing of cells within a colony geometry may be due to either their non-spherical character or their motility **[Schnyder et al. 2020]**.

### Equations for semi-confluent cell growth in 2D – circular colonies

Frequently cells grow as a monolayer, a situation resulting in 2D colony formation **[Martz and Steinberg, 1972; Wakita et al. 1994]**. In this work we consider colony growth in 2D to be isotropic, allowing us to equate colony radius with the number of cells within it via Eqn. 6 (where *r*_*culture*_ is the radius of the culture),

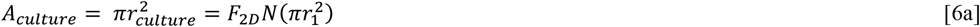

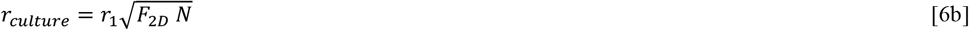

The number of cells within the perimeter region of radial thickness r_1_ is given by Eqn. 7.

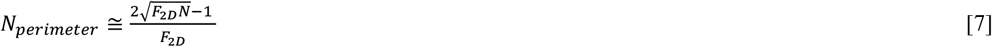

Substitution of Eqn. 7 into Eqn. 4 and its subsequent reinsertion into Eqn. 2 yields Eqn. 8a. This equation can be solved either by numerical integration or integrated directly between the limits [t=0, t] and [N(t=0)=1, N(t)] to produce Eqn. 8b.

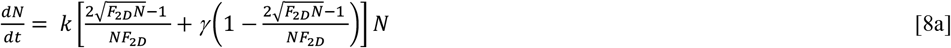

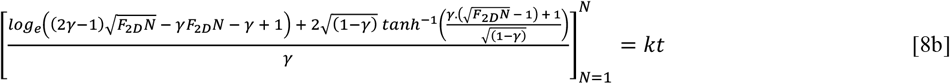

### Equations for semi-confluent cell growth in 3D – Spherical colonies

Cell culture can be carried out via hanging drop procedures **[Tung et al. 2011; Foty 2011; Zhao et al. 2019]** or within 3D culture environments **[Alessandri et al. 2013; Gunti et al. 2021]** to produce colonies having an additional dimensional aspect. One frequently observed 3D culture shape is a sphere, such as in the case of organoid development **[Karolak et al. 2018; Gunti et al. 2021]**. Spherical collections of cells are also seen in cancerous tumor development observed *in situ* in afflicted patients and in *in vitro* and *in vivo*^3^ laboratory experiments on cancerous growth **[Zietarska et al. 2007; de Lázaro et al 2021]**. The volume of a spherical colony is related to the number of cells within it by Eqn. 9,

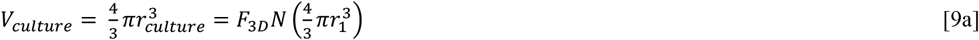

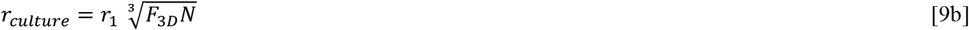

The number of cells growing within the perimeter regions for a spherical culture is given as Eqn. 10.

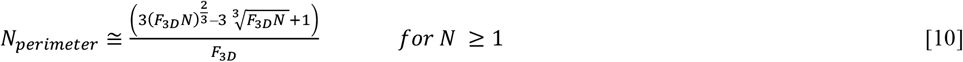

Substitution of Eqn. 10 into Eqn. 4 and its subsequent reinsertion into Eqn. 2 yields Eqn. 11a. This equation can be solved either by numerical integration or integrated directly between the limits [t=0, t] and [N(t=0)=1, N(t)] to produce Eqn. 11b.

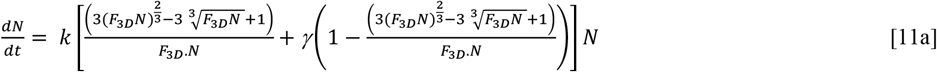

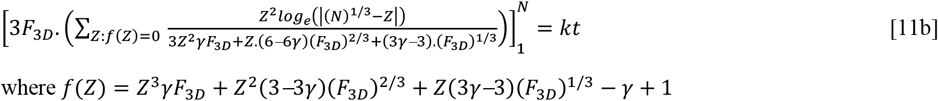

The summation in Eqn. 11b is carried out over the total number of solutions of *f(Z)* = 0. Having developed solutions for different forms of colony growth capable of exhibiting a variable level of local contact inhibition (indicated by values of β varying from 0 (minimum contact inhibition) to 1 (maximum contact inhibition)) we now show how such contact inhibition affects cell growth within colonies. The analytical equations describing colony growth (circular – Eqn. 8b; spherical – Eqn. 11b) are implicit in nature requiring that the solutions be determined by an iterative method where the total number of cells is fractionally altered until the LHS of the equation is equal to the RHS defined by the product of the intrinsic rate constant and the time **[Press et al. 2002]. Fig. 2** describes the rate of cell growth within the two colony types for five different levels of contact inhibition defined by values of γ = 1, 0.8, 0.5, 0.1 and 0.01 (with corresponding values of β = 0, 0.2, 0.5, 0.9 and 0.99). In interpreting Fig. 2 it is important to realize that all cells are characterized by an identical value of the intrinsic cell division rate k. For each case we note that increasing the extent of contact inhibition slows the rate of colony size increase. Significant effects on the colony growth rate due to differences in colony shape, are seen at greater extents of local contact inhibition.

**Figure 2:**
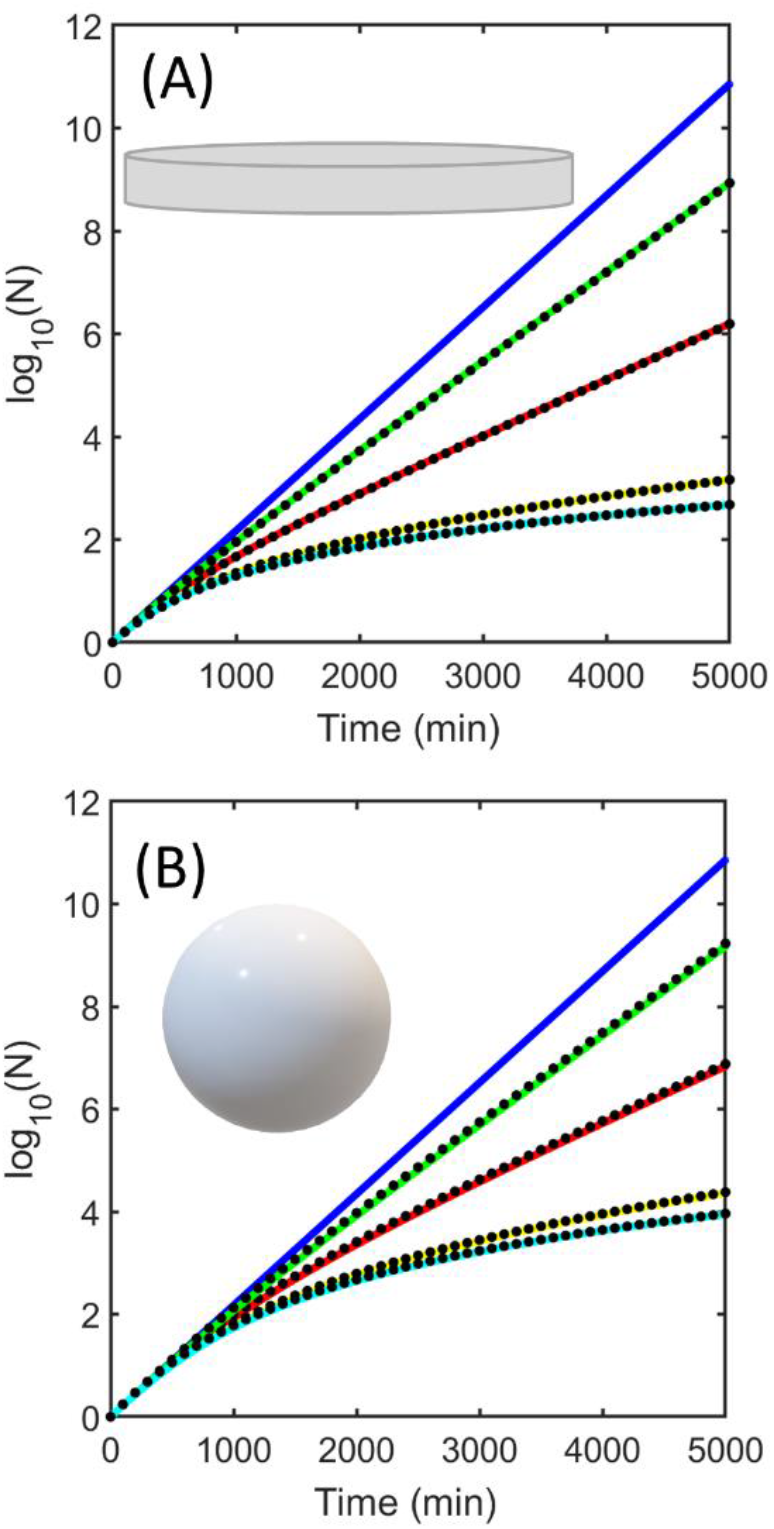
Effects of both geometry and extent of local contact inhibition on cell growth for cell growth in. (A) 2D monolayer and (B) 3D spherical colony. Simulation parameters k = 0.005 min−1, γ = 1 (blue), γ = 0.8 (green), γ = 0.5 (red), γ = 0.1 (yellow) and γ = 0.01 (cyan). Solid colored lines describe the numerical solution of the differential equations, black circles are the solution of the analytical equations.

## Discussion

The growth of cells within a collective is a fundamental tenet of biology that is central to topics such as, colony formation **[Wakita et al. 1994; Moore et al. 1973; Matsushita et al. 2004]**, the transition from unicellular to multicellular organisms **[Ros-Rocha et al. 2021]**, the growth of cancerous tumors **[Kapałczyńska et al. 2018; Gunti et al. 2021]**, and the creation of specialized tissues and organs **[Lewis-Israeli et al. 2021]**. When cell culture models are interrogated using techniques such as optical microscopy **[Wartenberg and Acker, 1995: Lawless et al. 2010; Chen et al. 2014; Hari et al. 2019]**, optical scanning densitometry **[Ruusuvuori et al. 2014]**, and atomic force microscopy **[Hoh et al. 1994; Gillis et al. 2012; Chatterjee et al. 2014]** the experimenter can access parameters able to inform upon the size and shape of the colony and the number of cells within it as function of position. However, interpretation of the rate of cell growth within a colony faces a number of complications. Aside from the question of contact inhibition (as treated here), the rate of cell growth may also be affected by differential nutrient transfer and depletion **[Burton, 1966; McElwain and Ponzo, 1977; Drasdo and Höhme, 2005; Banwarth-Kuhn et al. 2020; Golden et al. 2022]**, cellular motility and heterogeneity **[Waclaw et al. 2015]**, subsequent cellular differentiation and/or cell death **[Welker et al. 2021]**, and possible physical interaction of externally positioned cells with the growth matrix **[Ahmad-Khalili and Ahmad, 2015]**. To include such complications the rate of cell colony growth has been modelled using either continuum dynamics approaches **[Montel et al. 2015; Radszuweit et al. 2009]** or stochastic particle dynamics **[Lowengrub et al. 2010; Van Liedekerke et al. 2015; Hall, 2022]**. However, prior to the development of complex models of cell growth and division that include such higher-order factors **[e.g. see Van Liedekerke et al. 2015; Montagud et al. 2021]**, the ability to test simpler models, constructed on the basis of just a few parameters, possesses reductive value in assessing, via direct regression, the extent to which these other factors play (or do not play) a role. The equations presented within the present work fulfil this simple comparison function in a manner not requiring significant computational effort^4^ and potentially offer a pathway to direct analysis of cell culture experiments^5^. The approach could, in principle, be modified to account for position dependence of the level of contact inhibition by specifying separate regions of the interior and assigning a specific value of β to the cells growing in those regions, and this will form the main subject of part II of this work to be published at a later date. Whilst fully acknowledging that more complex models of cells growing in culture exist, it is hoped that the present equations may be used in a kind of Occam’s razor-type approach, to see if simple models based on a low parameter conceptualization of local contact inhibition, in terms of k and β/γ, may suitably describe the observed data, be potentially characteristic in nature and also be transferable across different cell culture experiments.

## Appendix One

The stated raison d’etre of the current paper was the development of a model that would be capable of describing and characterizing variable extents of contact inhibition affected cell growth using just one or two mechanistic parameters (in this case the cell growth rate k and the contact inhibition parameter γ). However, a fair critique of this work might involve asking for consideration of how this simple rationalization of contact inhibition compares against more complex models of cell growth. This criticism is partially answered within the following appendix.

As discussed within the main text, modelling of cell growth and division is complicated and any model will always be a gross approximation for reasons relating to the following.

i. A cell is not a uniform entity (e.g. the term cell can refer to eukaryotic, prokaryotic, archae forms and within multicellular organisms various differentiated types of cells exist or may manifest over time).
ii. All descriptions of cell growth and division are phenomenological i.e. we assign states based on a certain observation methodology (e.g. this can take the form of colony size, cell size and number, a particular marker of protein/nucleic acid expression, chemical uptake and chemical release).
iii. Cells respond to their environment (e.g. as open systems cells are affected by their local surroundings making each cell a unique entitity).

In wrestling with the above limitations quantitative biologists generate models of sufficient complexity to describe and analyze particular phenomena. The model developed in this paper sought to describe the most common observable – the size of a spherical or circular colony growth. The single parameter model used to describe unrestricted cell growth (Eqns. 1-3, Fig. 1) is perhaps the most fundamental of all cell growth models **[Robertson, 1923; Bell and Anderson, 1967]**. A well-posed two-compartment approximation was used to adapt this model for the case of variable contact inhibition and equations were developed therefrom. To extend the complexity of our approach here we introduce two slightly more sophisticated variations of cell growth corresponding to the cases of cell fission (in which a mature cell divides in two after which the two smaller newly divided cells then grow to mature form – Eqn. A1 and App. Fig. 1a) and cell budding (in which a mature cell initates the growth and eventual detachment of a small ‘bud’ daughter cell from its surface with the bud subsequently growing into a mature cell – Eqn. A2 and App. Fig. 1b) **[Balasubramnian et al. 2004]**.

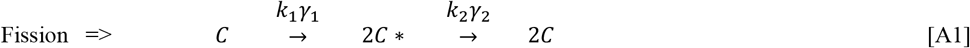

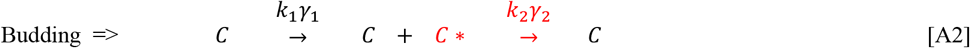

In these minimal representations each type of cell growth process (fission or budding) is defined by two rate parameters, k_1_ and k_2_, with another two parameters, γ_1_ and γ_2_, used to describe how each of these steps are affected by cellular contact inhibition. In a manner corresponding to the development of Eqns. 4-8 and Eqns. 9-11 we can define the number of cells of type C and C* as N_C_ and N_C*_, their average spherical radius as r_c_ and r_c*_, and the average space taken up by each cell type within a 2D colony in relation to their actual size as, F_2D_ and F_2D*_, and within a 3D colony as F_3D_ and F_3D*_. Proceeding with the case for 2D colony growth^6^ we use these terms to specify the geometry relations relevant to contact inhibited growth (Eqns A3a-d) where Ω is a nominated edge distance in which the contact inhibition effect is absent.

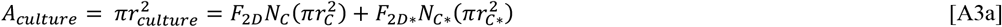

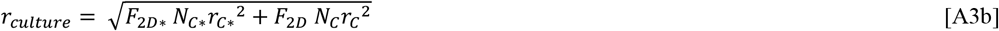

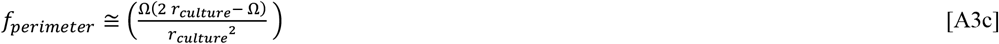

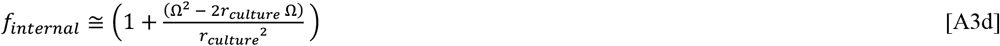

The rate equations defining partially contact inhibited cell growth (for the C and C* forms) in two dimensions for a fission mechanism can be written as equation set A4.

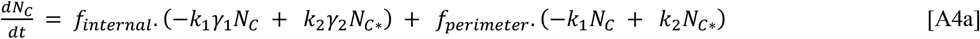

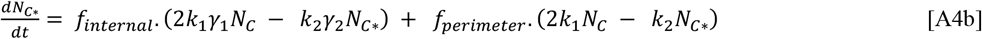

The corresponding rate equations defining partially contact inhibited cell growth (for the C and C* forms) in two dimensions for a budding mechanism can be written as equation set A5.

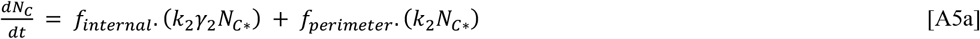

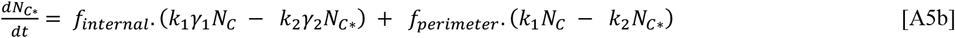

By adopting the two compartment approximation (internal and perimeter regions) used in the main paper for defining variable contact inhibition and implementing a number of extra parameters (Ω, k_1_, k_2_, γ_1_, γ_2_, F_2D_, F_2D*_) we note that a similar mathematical development can be arrived at – albeit only as a set of inter-related ordinary differential equations to be solved via numerical integration. Although possessing a slightly more realistic biological mechanism the development for fission and budding mechanisms shown in this appendix suffers from uncertainty in the assignment of parameters e.g. how differentially affected are the k_1_ and k_2_ steps by contact inhibition γ_1_ and γ_2_, and when considering an array of differently sized cells what is the correct value of Ω (or is this particular for different cell states)? In adopting a higher parameter model the transition from a dual purpose analysis/simulation role to a simulation only role is nearly fully effected. Within the simulation only domain, more complete and more complex models (of the stochastic agent type or continuous partial differential equation type) may prove to be of more benefit – we shall see in future.

## One Figures Appendix

**App. Figure 1:**
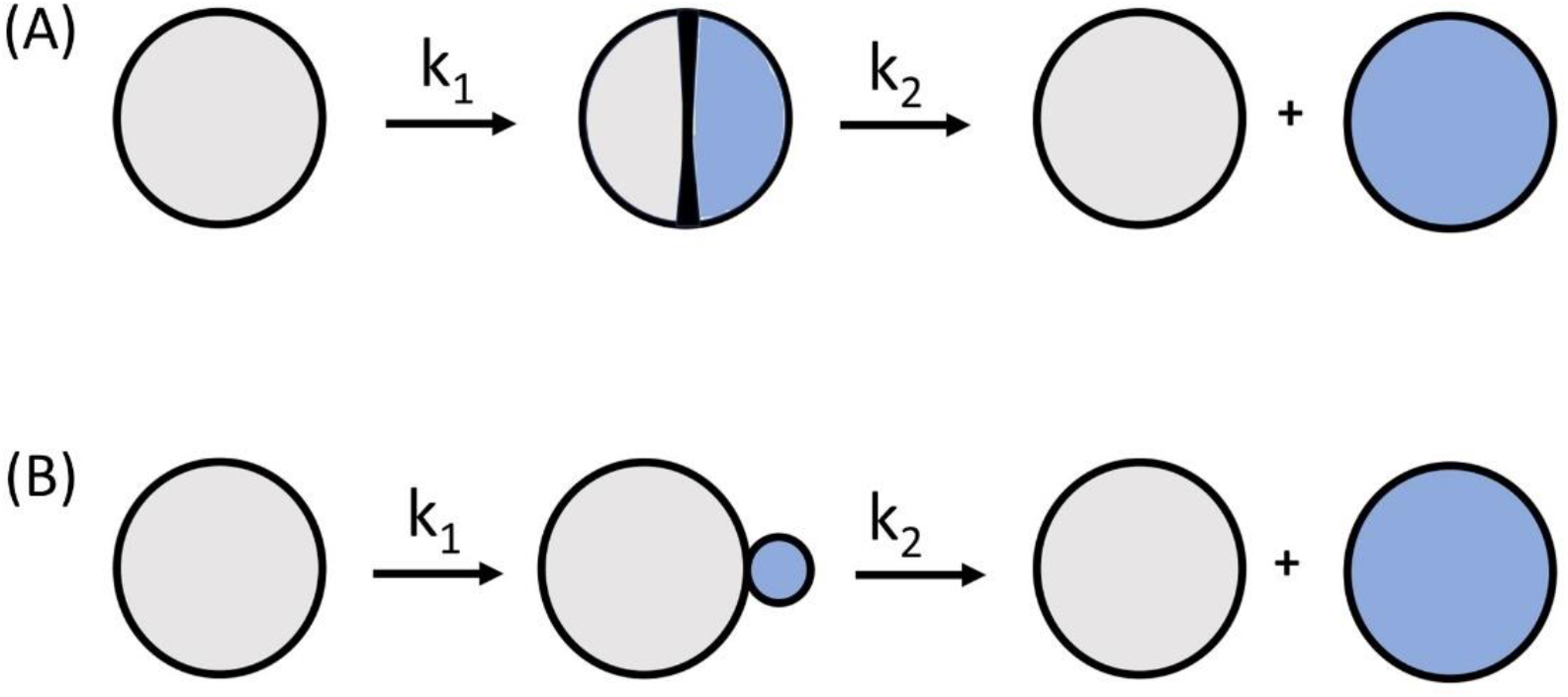
More complex models of cellular reproduction. (A) Fission type cell division in which the cell initially divides in two before then going on to regain a mature size (Eqns. A4). (B) Budding type cell division in which the cell initially produces a small daughter bud after which the bud grows to maturity (Eqns. A5).

## Acknowledgements

DH acknowledges helpful comments on an early version of this manuscript from Profs. Haruki Nakamura, Akira Kinjo and Kiyoshi Ohnuma. The author also acknowledges funding associated with the receipt of a ‘Tokunin’ Assistant Professorship carried out at the WPI-Center for Nano Life Science, Kanazawa University. This work was also supported, in part, by KAKENHI Start-Up grant 21K20633, KAKENHI Kiban-C grant 23K05712 and WPI ‘Funds in Aid of Research’ grant awarded to D.H.

## Footnotes

1 Which corresponds to the growth of cells in a well stirred liquid medium.

2 When contact inhibition is mediated by cell-to-cell chemical sensing this assumption is relatively strong. When contact inhibition is mediated by the force required to ‘push’ surrounding cells out of the way then the level of contact inhibition may be related to the position of the cell within the growing colony. We will address this latter case within the discussion section.

3 Such as for the case of xenograft or allograft tumor transplantation studies **[Sutherland et al. 1977; Zietarska et al. 2007]**.

4 Such as is the case for agent-based implementation of the Cellular Potts Model **[Van Liedekerke et al. 2020]**.

5 A similar type of two-state consideration of the differential form has been offered previously based on a more flexible definition of the interior and edge regions **[Mayneord, 1932; Montel et al. 2015; Radszuweit et al. 2009]**. These developments either remained in differential form or proceeded in a less straightforward way involving specification of a perimeter edge width and which assumed a step function to contact inhibited growth (i.e. it lacked the variable contact inhibition parameter γ). All approaches lacked the packing consideration that would allow for accommodation of asymmetry in the cultured cell type. Finally, the definition of contact inhibition employed in the present work ensures that the differential and integrated equations developed here are applicable to an initial condition corresponding to a single cell.

6 Allowing the reader to develop the 3D case as an exercise.

